# Fish-released kairomones affect *Culiseta longiareolata* oviposition and larval life history

**DOI:** 10.1101/2020.12.22.423951

**Authors:** Alon Silberbush

**Affiliations:** Department of Biology and the Environment, Faculty of Natural Sciences, University of Haifa, Israel

**Keywords:** Predator-released kairomones, life history, oviposition habitat selection

## Abstract

Several species of mosquitoes respond to the presence of kairomones released by larval predators during oviposition habitat selection and larval development. These responses may differ among mosquito species and do not always correlate with larval survival. This study examined the responses of the mosquito *Culiseta longiareolata* Macquart to kairomones released by three species of fish during oviposition, *Gambusia affinis* Baird and Girard, *Aphanius mento* Heckel and *Garra rufa* Heckel. In addition, the study examined the effects of kairomones released by *G. affinis* on larval development. Results show that ovipositing female avoided cues from larvivorous, but not algivorous fish. In addition, developing larvae metamorphosed slower and showed increased mortality when exposed to fish-released kairomones. Results suggest that the responses of this mosquito species to fish-released kairomones may be explained by its competitive ability.

## Introduction

Semiochemicals indicating the presence of predators are often an important part of predator detection mechanisms in addition to visual, audio and tactical cues (Kats and Dill 1998). Aquatic organisms often rely specifically on kairomones or other predator associated semiochemicals, such as alarm or disturbance cues to detect the presence of aquatic predators, since chemical cues travel better in water in comparison to other cue types (Dodson et al. 1994, Chivers and Smith 1998, Weiss et al. 2012, Weissburg 2012). Aquatic organisms respond to predator-released kairomones in numerous ways including altered foraging behavior (Bucciarelli and Kats 2015), morphological changes such as the appearance of spikes in Crustaceans (Weiss et al. 2016), or color variations (McCollum and Leimberger 1997). Species with complex life histories, such as amphibians and aquatic insects spend their juvenile life stages in water, while terrestrial adults oviposit, or larvaeposit, in selected aquatic sites. Those species often rely on predator-released kairomones to detect larval predators when selecting a suitable habitat for their offspring (Blaustein 1999).

Ovipositing mosquito female detect and avoid water sources that contain predator-released kairomones originating from invertebrate larval predators (Chesson 1984, Silberbush et al. 2010, Ohba et al. 2012) and larvivorous fish (Angelon and Petranka 2002, Van Dam and Walton 2008, Eveland et al. 2016). Fish-released kairomones also affect larval behavior in addition to time and size at metamorphosis (van Uitregt et al. 2012, Roberts 2014).

The responses of mosquitoes to fish-released kairomones are often not straightforward. For example, ovipositing *Culex* female are strongly repelled by fish-released kairomones (Angelon and Petranka 2002). However, this repellency response is limited only to some species of fish, while others are ignored, and thus does not match the threat to larvae (Eveland et al. 2016, Segev et al. 2017, Silberbush and Resetarits Jr. 2017, Cohen and Silberbush 2020). Mosquito larvae developing in fish-conditioned water often reduce foraging activity (van Uitregt et al. 2012, Roberts 2014). In the overwhelming majority of case studies with larval amphibians and aquatic insects exposed to predator-released kairomones, this behavior is coupled with delayed time to metamorphosis (Relyea 2007). *Culex* larvae, on the other hand, respond to fish released kairomones by accelerating time to metamorphosis (Silberbush et al. 2015). This response is shown towards several species of fish-released kairomones (Cohen and Silberbush 2020), takes place despite reduction of foraging activity and does not depend on nutrient availability (Silberbush et al. 2019).

This study examined the responses of ovipositing *Culiseta longiareolata* Macquart (Diptera: Culicidae) to kairomones released by three fish species. The invasive-larvivorous western mosquitofish *Gambusia affinis* Baird and Girard, the native-larvivorous iridescent tooth carp *Aphanius mento* Heckel and the native-algivorous red garra, *Garra rufa* Heckel. In addition, the study examines the effects of Gambusia-released kairomones on larval development. This mosquito species is extremely abundant in the Middle East, Europe and Africa (Van Pletzen and Van der Linde 1981). Female oviposit, and larvae dwell, in temporary freshwater pools that are often inhabited by the common *Culex* species such as *Culex pipiens* Linnaeus and *C. laticinctus* Edwards (Margalit and Tahori 1974). Similar to *Culex* species, *C. longiareolata* female oviposit egg-rafts rather than single eggs. The probability of a specific female to survive more than one gonotrophic cycle is low (Spencer et al. 2002). *Culiseta longiareolata* female are repelled by predator-released kairomones during oviposition (Silberbush et al. 2010, Alcalay et al. 2019). Larval *C. longiareolata* were also shown to reduce foraging activity in response to fish-released kairomones (Roberts 2014). It is therefore hypothesized that ovipositing female and developing larval *C. longiareolata* will show a similar response to fish-released kairomones as *Culex* species who share a similar larval environment.

## Materials and methods

### Oviposition

A field experiment was set at Oranim campus Tivon, Israel, using black plastic tubs containing approximately 51L (66.04×50.8×15.24 cm) as oviposition pools. Pools were filled with aged tap water and 10g of rodent pellets (Ribos rodent chow-17% protein) were added to each pool in order to encourage oviposition. Fish were placed in plastic cylindrical cages (22 cm diameter) with multiple (1 mm diameter) holes and openings covered with a fiberglass screen. Pools were placed in sets of three, and treatments (fish species or fishless control) were randomly assigned within each set of three adjacent pools (block). Pools were checked and egg rafts removed daily. Larvae were identified to species after developing to fourth instar. The experiment was run twice. The first iteration, 21 March -15 May, with pools containing *G. affinis* and *A. mento*, the second from 8 April-May 1 the following year with pools containing *G. affinis* and *G. rufa. Aphanius mento* were collected from the breeding center at the botanical garden at Oranim campus, and the other two species from nearby sites. Individuals of all three species were of similar size (∼2-5 gr), fish biomass within a block was matched as much as possible. Fish were kept in 1200-L holding tanks and rotated every three days.

### Larval life history

Eighteen *C. longiareolata* egg rafts were collected from temporary pools around Oranim Campus. Egg rafts were hatched individually in aged tap water and the sibling larvae were randomly distributed among the two treatments, described below within a block, 24 hours from hatching. Thirty 1^st^ instars were placed in cups filled with 400 ml of aged tap water. Cups were kept at 25.7±3.2 °C (mean ±SD) and 14:10 light: dark photoperiod. Water in each cup was replaced daily and larvae fed with 0.1g of a finely ground mixture of Sera fish food (42.2% crude protein) and Ribos rodent chow (17% protein). These densities and food amounts were considered to be ideal for larval development (Van Pletzen and Van der Linde 1981, Al-Jaran and Katbeh-Bader 2001). Pupae were counted daily, removed and kept in separate vials until adult emergence, when they were classified by sex.

#### Fish conditioned water

Each block was comprised of two cups, fish conditioned water and a fishless control cup. Fish conditioned water were taken from 3 l tubs filled with aged tap water containing 2 *G. affinis* fish. Fish in the tubs were not fed but rotated every 3 days among a larger container where they were fed with the same Sera fish food given to the larvae. Fishless control cups were filled from similar tubs without fish.

### Statistical analysis

#### Oviposition

The number of egg rafts deposited by *C. longiareolata* females was summed, per pool, across all dates. The square root values of the summed values were used to improve homogeneity of variance, which was tested using Levene’s test. A value of 0.5 was added to the summed value 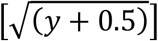 to stabilize population variance and meet the assumptions of Analysis of Variance (Yamamura 1999). Treatment means were compared using Fisher’s Protected LSD only when the main effect of treatment had p< 0.10, using α = 0.05 for individual LSD comparisons.

#### Larval development and survival

The average, per cup, number of days to pupation was used as a measure of larval development time, and the number of emerging adults per cup as measurement of survival. The square root values were used to improve homogeneity of variance of number of days to pupation, which was tested using Levene’s test. Data was analyzed using splitplot ANOVA, with sex (male and female) as the within-plot (cup) factor and treatment (fish-conditioned water or fishless control) as a fixed factor. Blocks (plot-egg raft) were used as a random factor. All analyses used SPSS statistics for windows version 24 (IBM Released 2016).

## Results

### Oviposition

The first experiment (*Gambusia* and *Aphanius*) received a total of 234 *C. longiareolata* egg rafts. Distribution varied significantly among treatments (F_2,10_=4.25; p=0.046) with the mean number of egg rafts in control pools significantly higher than in both fish-containing pools (figure 1a). In the second experiment (*Gambusia* and *Garra*) a total of 139 *C. longiareolata* egg rafts were found. Distribution varied significantly among treatments (F_2,10_=3.36; p=0.077) with the mean number of egg rafts in control pools significantly higher than in *Gambusia* pools, but not from egg rafts oviposited in *Garra* treated pools (figure 1b).

**Figure 1.**
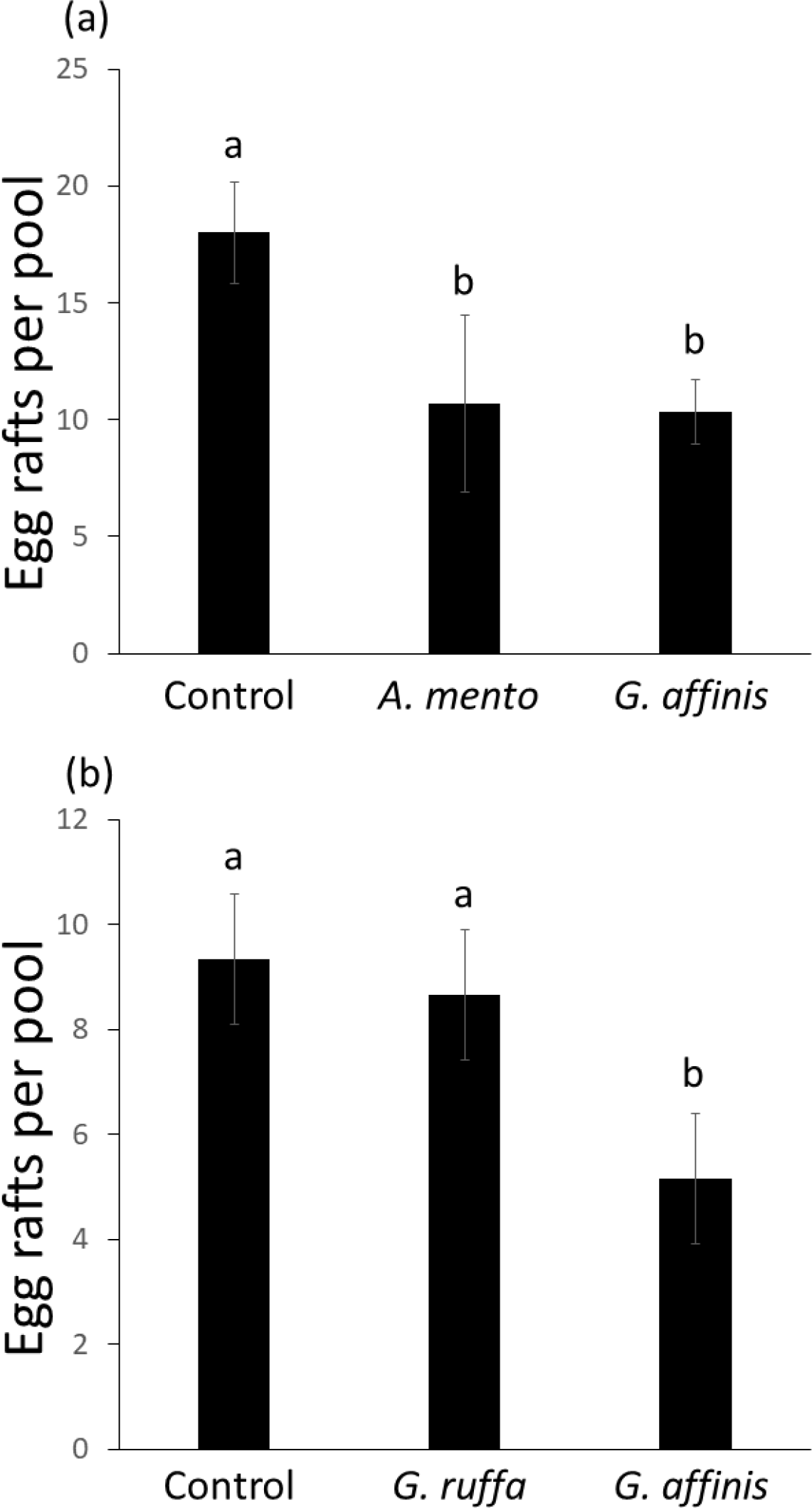
Oviposition of *Culiseta longiareolata* in the two field experiments: (a) *Aphanius* and *Gambusia*; (b) *Garra* and *Gambusia*. Numbers are mean number of egg rafts per pool (±1 SE). Letters indicate treatments that are significantly different.

### Larval development

A total of 858 larvae emerged as adults (79.4% of the original 1^st^ instars). Significantly more females emerged than males per cup (sex effect: F_1:17_=16.99; p=0.001). This sex ratio bias was not dependent on fish kairomones (sex X treatment interaction: F_1:17_=0.14; p=0.71). An average ±SD of 3.9± 6.4 more larvae survived to adulthood in fishless control cups in comparison to cups with fish-conditioned water (treatment effect: F_1:17_=6.56; p=0.02; figure 2a).

**Figure 2.**
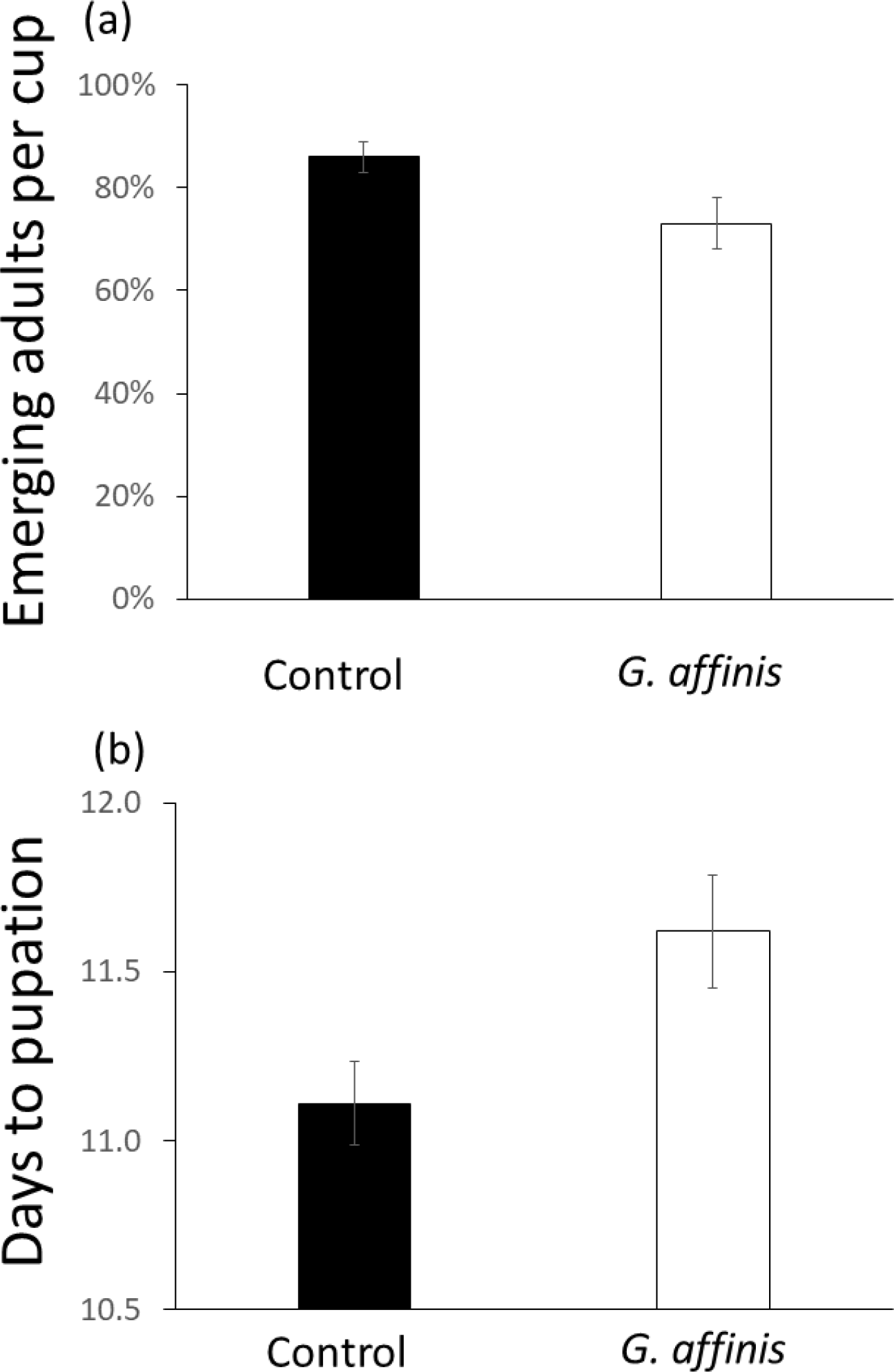
Larval survival (a) and time to pupation (b) in cups containing *Gambusia*-conditioned or fishless-control water. Numbers are respectively mean larval survival and mean time to pupation per cup (±1 SE). Bar colors indicate treatments that are significantly different.

### Days to pupation

males pupated faster than females (sex effect: F_1:17_=168.05; p<0.001), with no significant sex X treatment effect (F_1:17_=0.02; p=0.88). Larvae exposed to fish-conditioned water pupated significantly slower by average ±SD of 0.55± 1.2 days in comparison to larvae in fishless-control cups (F_1:17_=5.5; p=0.03, figure 2b).

## Discussion

In theory, prey’s response to predator-released kairomones should be proportional to the level of predation risk (Weiss et al. 2012, Wisenden 2015). While behavioral shifts can be affective against general threats, more expensive responses such as life history alternations, are assumed to be implied against more immediate dangers (Weiss et al. 2012).

The aim of this study was to show the responses of *C. longiareolata* to fish-released kairomones representing two levels of risk. Ovipositing adult female facing threat to future offspring, and developing larvae who face a more immediate risk. Results show that ovipositing female were strongly repelled by kairomones of both native and invasive larvivorous fish but not the algivorous *Garra rufa* (figure 1). This response points to the ability of females to distinguish between kairomones released by different fish species, and is also directly coordinated with the level of threat to future larval offspring. The original hypothesis suggested that the response of *C. longiareolata* to fish-released kairomones should be similar to that of the *Culex* species who share similar larval habitats. This hypothesis was not met since the responses of *Culex* species to fish-released kairomones during oviposition habitat selection do not always match the threat to larvae. While aquatic insects and amphibians avoid most species of fish during oviposition (Resetarits Jr. and Binckley 2013), several species of *Culex* exhibit a more specific reaction. For example, ovipositing *Culex restuans* and hybrid *C. pipiens*-*quinquefasiatus* (*C. pXq*) were both repelled by kairomones released by larvivorous mosquitofish *G. affinis* in outdoor mesocosms. This repellent effect was not shown with response to green sunfish, *Lepomis cyanellus*, or the pirate perch, *Aphredoderus sayanus* (Eveland et al. 2016, Silberbush and Resetarits Jr. 2017). Similar species-specific responses to fish-released kairomones were recorded in five other species of *Culex* associated with breeding in large bodies of water (Segev et al. 2017, Cohen and Silberbush 2020). *Culiseta longiareolata* are considered early colonizers of newly flooded pools (Blaustein and Margalit 1994, Ward and Blaustein 1994, Spencer et al. 2002). As such, they are less likely to encounter larvivorous fish. Avoidance of larvivorous fish is not always shown by mosquito species who typically oviposit in small or temporary bodies of water that are not likely to contain fish (Van Dam and Walton 2008, Walton et al. 2009). The general avoidance response of ovipositing *C. longiareolata* may point toward a larger aquatic niche of this species.

Aquatic larvae developing under the threat of predation are hypothesized to exhibit stress-associated symptoms such as reduced survival (Relyea 2007). In addition, predator presence leads to reduced foraging which is hypothesized to result in an extended immature period due to reduced consumption of nutrients (Werner 1986, Ludwig and Rowe 1990, Rowe and Ludwig 1991). Larvae in this study exhibited a ∼13% reduction in survival to adulthood in addition to delayed metamorphosis, when exposed to fish-released kairomones (figure 2). Further study is needed to examine whether rapidly developing larvae within a cohort, are also more likely to show a stronger response to stress-related reactions. However, both reduced survival and longer immature periods are the most widespread responses of aquatic larvae developing under threat of predation (Relyea 2007). As before, the original hypothesis, assuming similarity to the response of *Culex* larvae exposed to fish-released kairomones was not met. Larval *Culex* are among the exceptional who respond to fish-released kairomones by accelerating time to metamorphosis with no recorded reduction in survival to adulthood (Silberbush et al. 2015, Silberbush et al. 2019, Cohen and Silberbush 2020).

Response to predator presence may depend on other factors as well. In theory, an ovipositing female may select a predator-containing pool to avoid competitive effects (Spencer et al. 2002). Indeed, a subordinate mosquito competitor exhibited this phenomenon in response to the presence of *C. longiareolata* larvae (Silberbush et al. 2014). By contrast, *C. longiareolata* female avoid ovipositing in predator-containing pools even when the alternative is a high density of conspecifics (Kiflawi et al. 2003). The dominance competitive abilities of this species along with its aggressive behavior (Blaustein and Margalit 1994, Tsurim et al. 2013) may therefore explain this general avoidance of larvivorous fish. Competition may also affect time to metamorphosis, larval *C. longiareolata* may reduce (Tsurim et al. 2013) or increase (Blaustein and Margalit 1994, Tsurim et al. 2013) this period depending on competitor’s identity.

In conclusion, results of this study point to the possibility that response to the presence of predators may also depend on the competitor abilities of the prey species.

## Acknowledgments

I wish to thank Sagiv Cohen for assistance in data collection. Hatem Abu Raiya and Alon Ornai provided logistical support. Avi Bar-Massada and Yoram Gerchman for fruitful conversations. There was no conflict of interest regarding this study.

